# Mapping Individual Differences in Intermodal Coupling in Neurodevelopment

**DOI:** 10.1101/2024.06.26.600817

**Authors:** Ruyi Pan, Sarah M. Weinstein, Danni Tu, Fengling Hu, Büşra Tanrıverdi, Rongqian Zhang, Simon N. Vandekar, Erica B. Baller, Ruben C. Gur, Raquel E. Gur, Aaron F. Alexander-Bloch, Theodore D. Satterthwaite, Jun Young Park

## Abstract

Within-individual coupling between measures of brain structure and function evolves in development and may underlie differential risk for neuropsychiatric disorders. Despite increasing interest in the development of structure-function relationships, rigorous methods to quantify and test individual differences in coupling remain nascent. In this article, we explore and address gaps in approaches for testing and spatially localizing individual differences in intermodal coupling, including a new method, called CEIDR. CEIDR controls false positives in individual differences in intermodal correlations that arise from mean and variance heterogeneity and improves statistical power by adopting adaptive cluster enhancement. Through a comparison across different approaches to testing individual differences in intermodal coupling, we delineate subtle differences in the hypotheses they test, which may ultimately lead researchers to arrive at different results. Finally, we illustrate these differences in two applications to brain development using data from the Philadelphia Neurodevelopmental Cohort.

## 1 Introduction

During adolescence, the developing brain undergoes a profound structural and functional reorganization. To understand how the brain changes during childhood, researchers increasingly use multimodal neuroimaging to measure both distinct and shared spatial patterns in features of brain anatomy and function. Measures of *intermodal coupling* have been increasingly explored in recent research (Avants et al. 2010; Devonshire et al. 2012; Ouyang et al. 2015; Phillips et al. 2016; Vandekar et al. 2016; Iadecola 2017; Baum et al. 2020; Baller et al. 2022; Hu et al. 2022). Unlike studying individual differences separately in different brain modalities, the use of intermodal coupling allows researchers to study relationships between brain features (e.g., structure-function) that are thought to be critical for mechanisms underlying risk for developmental psychopathology, cognition, and brain health across the lifespan.

Despite the emerging utility of intermodal coupling measures in developmental cognitive neuroscience research, methods for conducting statistical testing on individual differences in coupling have yet to be assessed in terms of their statistical power (i.e., replicability) and interpretability. Also, to the best of our knowledge, no clear generative model currently exists that parametrizes intermodal coupling and its individual differences, which may introduce unwanted variability in results due to limitations in the analysis methods. Amid growing concerns about reproducibility in neuroimaging-based studies of neurodevelopment (Flournoy et al. 2020; Klapwijk et al. 2021; Marek et al. 2022; Botvinik-Nezer et al. 2020), this article seeks to concretize both the formulation and evaluation of hypotheses about intermodal coupling. To this end, we use the lens of statistical hypothesis testing to formulate scientific questions about individual differences in intermodal coupling. Even in the presence of confounding and other sources of noise in imaging data, such tests can be useful for identifying both *if* and *where* individual differences in coupling exist.

To illustrate the broad applicability of our method, we leverage data from a large neuroimaging cohort to investigate how intermodal coupling evolves in the course of healthy brain development. Specifically, we first focus on intermodal coupling between cortical thickness (CT) and sulcal depth (SD), measured using structural MRI. The overall volume of the cortex generally increases in early childhood, then peaks and decreases during late childhood and adolescence (Raznahan et al. 2011). This decrease in volume is mainly driven by cortical thinning that preserves surface area (Kelly et al. 2023), although thinning does not occur uniformly across the brain, with some prior work demonstrating dissociable processes of thinning in the sulci but thickening in gyri (Vandekar et al. 2016). Therefore, a high-resolution characterization of the relationship between cortical thickness and cerebral surface anatomy may shed light on the mechanisms supporting neurodevelopment.

To demonstrate the generalizability of our method across feature types, we also evaluate whether the relationship between amplitude of low-frequency fluctuations (ALFF) and cerebral blood flow (CBF). From blood oxygen level dependent (BOLD) functional magnetic resonance imaging (fMRI), we use ALFF signals as a proxy of neuronal activity; from arterial spin labelling, we can quantify neurovascular perfusion through CBF. Thus, for each individual, each location (e.g., voxel or vertex) in the brain image is associated with a value of CBF and ALFF, yielding two brain maps. Prior imaging studies focusing on CBF and ALFF have revealed age-related changes in neurologically healthy adolescents (Satterthwaite et al. 2014; Taki et al. 2011), as well as changes between healthy adults and people with cognitive impairment (Chen 2019; Hoptman et al. 2010), suggesting that both blood oxygenation and perfusion are critical for executive function. While CBF and ALFF brain maps offer valuable insights into brain function individually, examining the coupling between them together provides a deeper understanding of neurovascular coupling. Recent research suggests that neurovascular coupling may support cognitive functioning (Stackhouse and Mishra 2021; Phillips et al. 2016); in particular, Baller et al. 2022 demonstrated that CBF-ALFF coupling may be linked to changes in executive function in healthy adolescents, and that this coupling was sex-, age-, and region-dependent.

The overarching goal of this article is to describe statistical approaches that may be used to quantify, test, and spatially localize individual differences in intermodal coupling. We then propose several extensions to existing methodology. First, we conceptualize the symmetric relationship between two imaging modalities as a conditional correlation (Yan and Fine 2004; Tu et al. 2024), which can vary by age, sex, or other variables of interest. Finally, we propose CEIDR (**<u>C</u>**luster **<u>E</u>**nhancement for testing **<u>I</u>**ndividual **<u>D</u>**ifferences in *ρ* (**<u>r</u>**)), a statistical inference procedure to test individual differences in (conditional) intermodal correlations, which we illustrate as achieving high power and spatial localization. We apply our method to multi-modal MRI data from the Philadelphia Neurodevelopmental Cohort (PNC) (Satterthwaite et al. 2014), delineating both robust developmental effects and sex differences in intermodal coupling.

## 2 Methods

### 2.1 Notation

For subject *i* = 1, …, *N* and location (i.e., vertex or voxel) *v* = 1, …, *V*, we denote brain measurements for two modalities as *y*_1*i*_(*v*) and *y*_2*i*_(*v*), respectively. And, we let *x*_*i*_ represent the covariate of interest and **z**_*i*_ represent the nuisance covariate vector being adjusted for subject *i*. Categorical variables could be dummy-coded to be used in statistical modeling in our paper.

### 2.2 Parametric model and the null hypothesis for conditional correlation

To clearly define subject-level intermodal coupling, we consider the following parametric model. For the *m*th imaging modality, we consider the following mean and variance functions depending on covariates (i.e., *µ*_*m*_(**z**, *x*_*i*_, *v*) and *σ*_*m*_(**z**_*i*_, *x*_*i*_, *v*)) for a model for *y*_*mi*_(*v*):

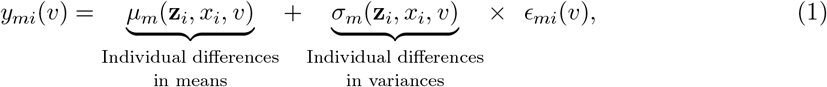

where the residual terms *ϵ*_1*i*_(*v*) and *ϵ*_2*i*_(*v*) follow the joint distribution given by:

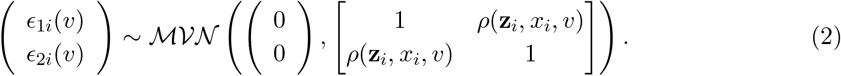

Here, the residual correlation *ρ*(**z**_*i*_, *x*_*i*_, *v*) is the subject-level intermodal coupling of our interest. We hypothesize that *ρ*(**z**_*i*_, *x*_*i*_, *v*) for subject *i* is a function of the variable of interest *x*_*i*_ (e.g., age) and other potential confounding variables **z**_*i*_ (e.g., sex), given by:

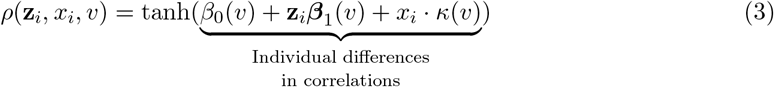

where tanh(*·*) is a hyperbolic tangent link function ensuring that the correlations lie strictly between −1 and 1 (Yan and Fine 2004; Tu et al. 2024).

Given our model formulations in Equations (1), (2), and (3), we consider the following null hypothesis:

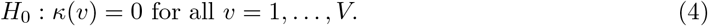

From the model, we note that individual differences between two imaging modalities may arise in the means *µ*_*m*_(**z**_*i*_, *x*_*i*_, *v*) and variances *σ*_*m*_(**z**_*i*_, *x*_*i*_, *v*) from each modality, as well as their ‘residual’ correlations *ρ*(**z**_*i*_, *x*_*i*_, *v*). Therefore, a nonzero *κ*(*v*) indicates that intermodal coupling at vertex *v* is associated with the variable of interest *x*_*i*_ after both (i) adjusting for covariate effects on mean and variance structures and (ii) adjusting for other nuisance covariate effects from **z**_*i*_ on residual correlations.

### 2.3 When could false positives in intermodal correlation be inflated?

As correlation naturally depends on mean and variance, the models for mean and variance should be correctly specified to accurately specify correlation models and to prevent possibly inflated false positives. We point out two possible misspecifications in mean and variances. First, the “observed covariate effects” on mean and variance could be misspecified when, for example, there are non-linear age effects on imaging data but non-flexible models (e.g., linear age effects) are assumed in Equation (1) (Fjell et al. 2009; Ziegler et al. 2012; Tamnes et al. 2013). A relevant case would be that an interaction between covariates is present but left out in the model. Second, it is possible that some important covariates are not collected in the study design and therefore cannot be included properly in Equation (1) (“unobserved covariate effects” on mean and variance).

### 2.4 CEIDR

We develop a powerful method to test the null hypothesis in Section 2.2, ensuring robust model specification, high sensitivity through cluster enhancement, and proper false positive control through permutation. We outline the five main stages of CEIDR below (See Figure 1 for illustrations).

**Figure 1.**
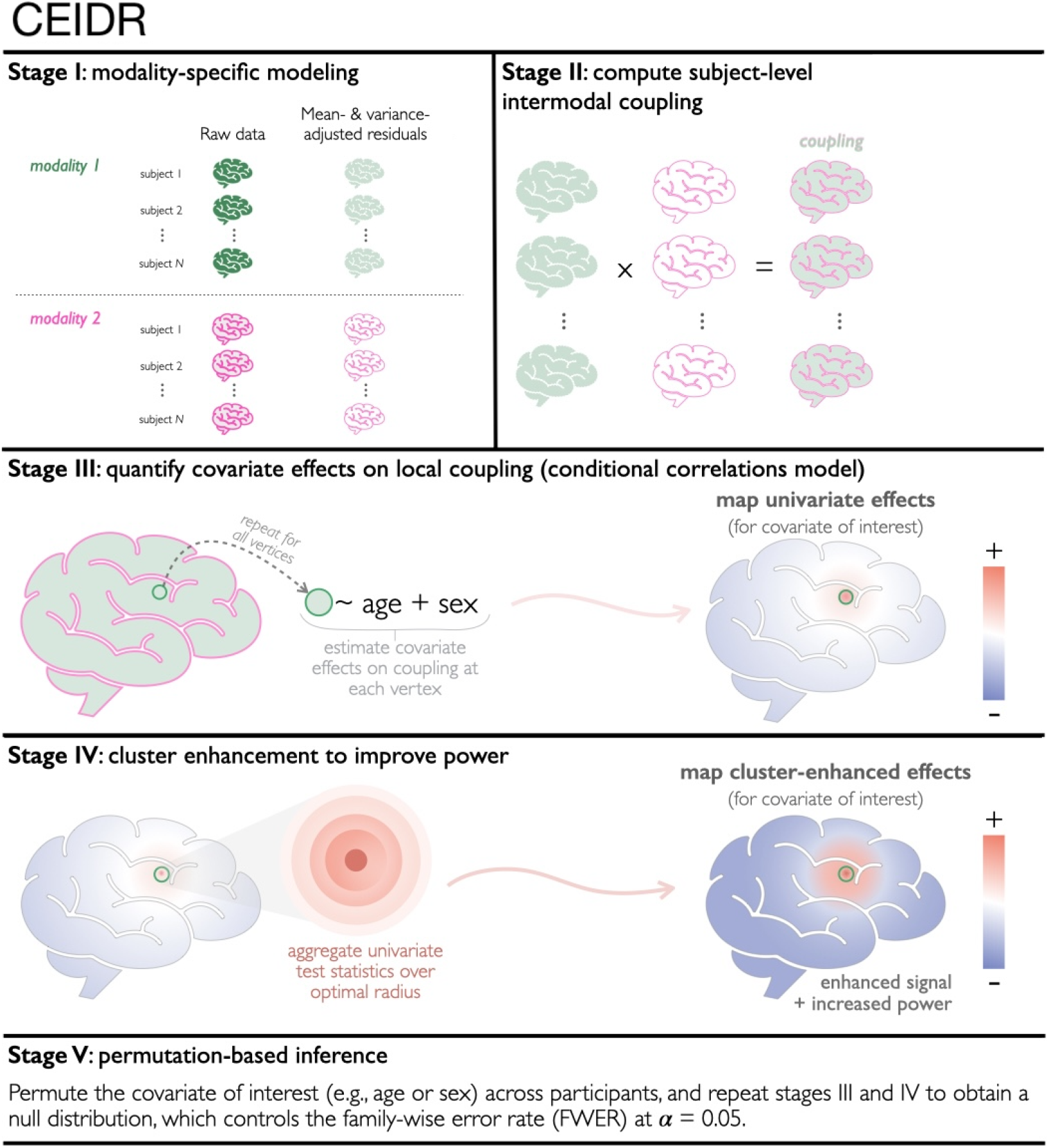
Overview of the proposed method, CEIDR.

I. *Removing unrelated data variability*. To mitigate positive findings that may be driven by unrelated variability in the marginal means and variances of each modality, we propose two approaches for performing marginal mean and variance adjustment.
  I.1 *Between-subject adjustment*. We use information from the full sample to adjust for co-variates like age and sex. Supported by the nonlinear brain structural dynamics found in prior neuroimaging studies (Ziegler et al. 2012; Tamnes et al. 2013), we fit the generalized additive model (GAM), which allows for flexible specification of both linear and nonlinear covariate effects, to estimate individual means *µ*_*m*_(**z**_*i*_, *x*_*i*_, *v*) and variances *σ*_*m*_(**z**_*i*_, *x*_*i*_, *v*) of each modality at each vertex. In this article, we consider *µ*_*m*_ to be modeled by a main effect of sex and a cubic-spline of age stratified by sex, and log(*σ*_*m*_) to be modeled by the main effect of age and sex, which can be specified in the `gamlss` package in R. Once *µ*_*m*_(**z**_*i*_, *x*_*i*_, *v*) and *σ*_*m*_(**z**_*i*_, *x*_*i*_, *v*) are estimated, we obtain the standardized residuals as follows:

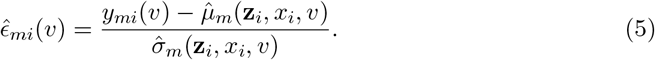
  I.2 *Within-subject adjustment*. One can also use local information within each subject to adjust the individual means and variances. For each subject, assuming local neighbors have the same means and variances, we use the sample mean and variance within a local region *N*_*r*_(*v*), i.e., the set of all vertices whose distances from the vertex *v* is less than or equal to *r*, to estimate the subject-specific mean and variance at vertex *v* and do adjustment:

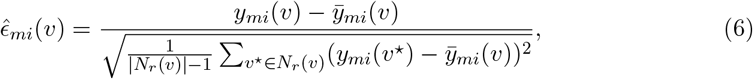

where 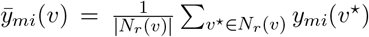. We selected a radius of *r* = 5mm as the default value in this article, which is suitable for achieving robust coupling estimations. The 5mm radius remains appropriate for this resolution, as most vertices with approximately 10 neighbors (the median of |*N*_5_(*v*)| across all vertices in fsaverage5 is 11). One can choose either approach depending on the study context. Additionally, the two strategies can be combined: first applying the between-subject adjustment using observed covariates, and then performing a within-subject adjustment (on 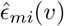 from Equation (5)) to account for potential unknown factors that may still influence the marginal mean and variance. In this paper, we use both between- and within-subject adjustments in our data analysis.
II. *Estimating subject-level intermodal coupling*. We compute subject-level measures of intermodal associations using the product of the modality-specific residuals:

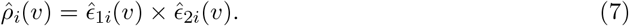
III. *Modeling* 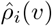 *in terms of covariates*. We assume the covariates regulate *ρ*_*i*_(*v*) through tanh(*·*) link function specified in Equation (3). Although tanh(*·*) is not straightforward to model using a GLM, we observe that tanh(*t*) *≈ t* when |*t*| is small or moderate (e.g., less than 0.5), which is consistent with the magnitude of correlations often found in neuroimaging studies (see Figures 3 and 4 in Section 3). Under this approximation, we model the relationship between intermodal coupling and covariates using a GLM formulation of

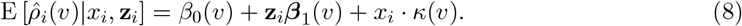

**Figure 2.**
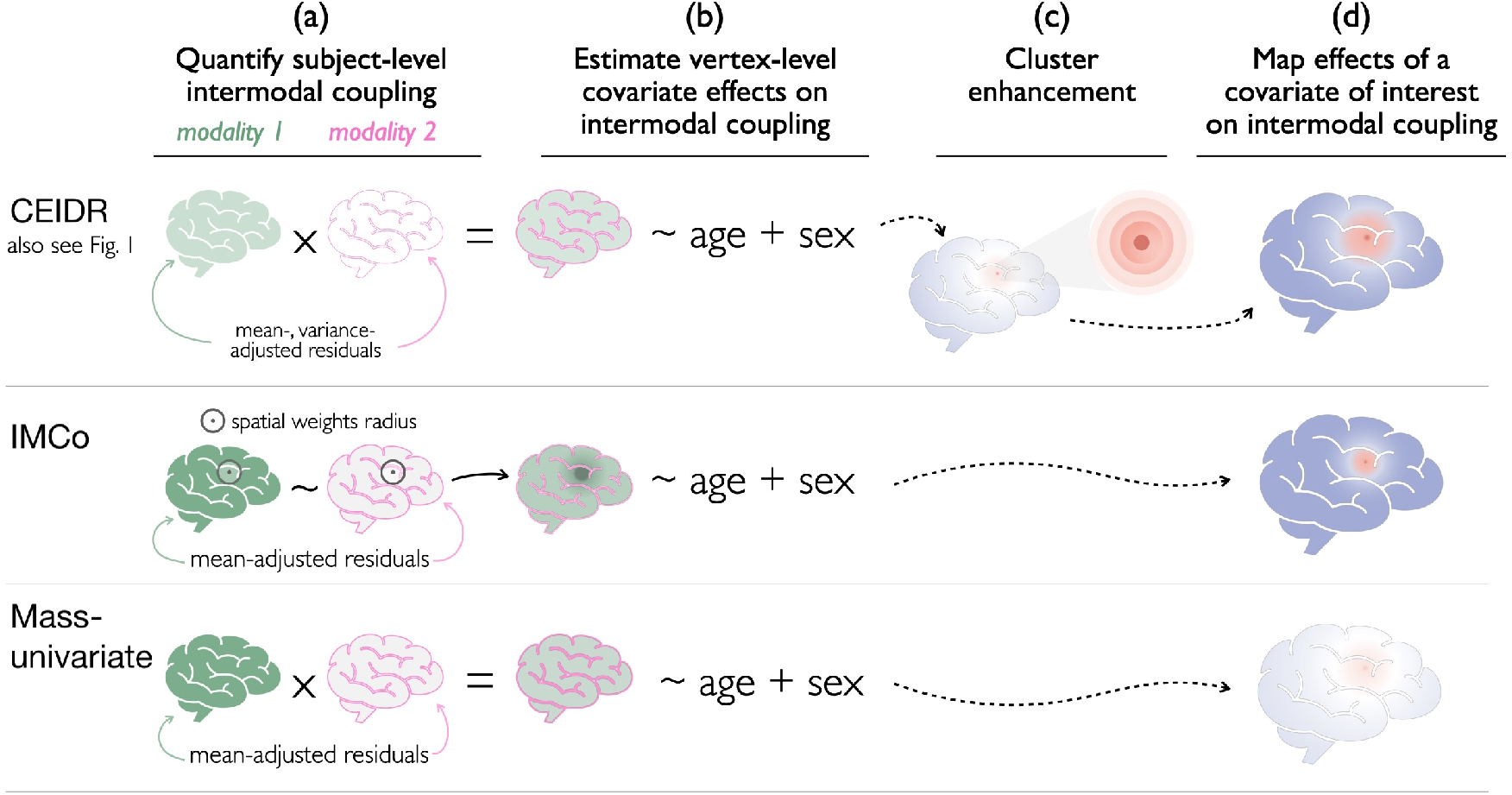
Overview of the proposed method, CEIDR, along with two alternative approaches to quantifying intermodal coupling (a-b) and individual differences (c-d). A more detailed illustration of CEIDR is provided in Figure 1. We perform statistical inference for all three methods using the same permutation-based procedure, which uses a brain-wide threshold for statistical significance in order to control the family-wise error rate (FWER) at *α* = 0.05.

**Figure 3.**
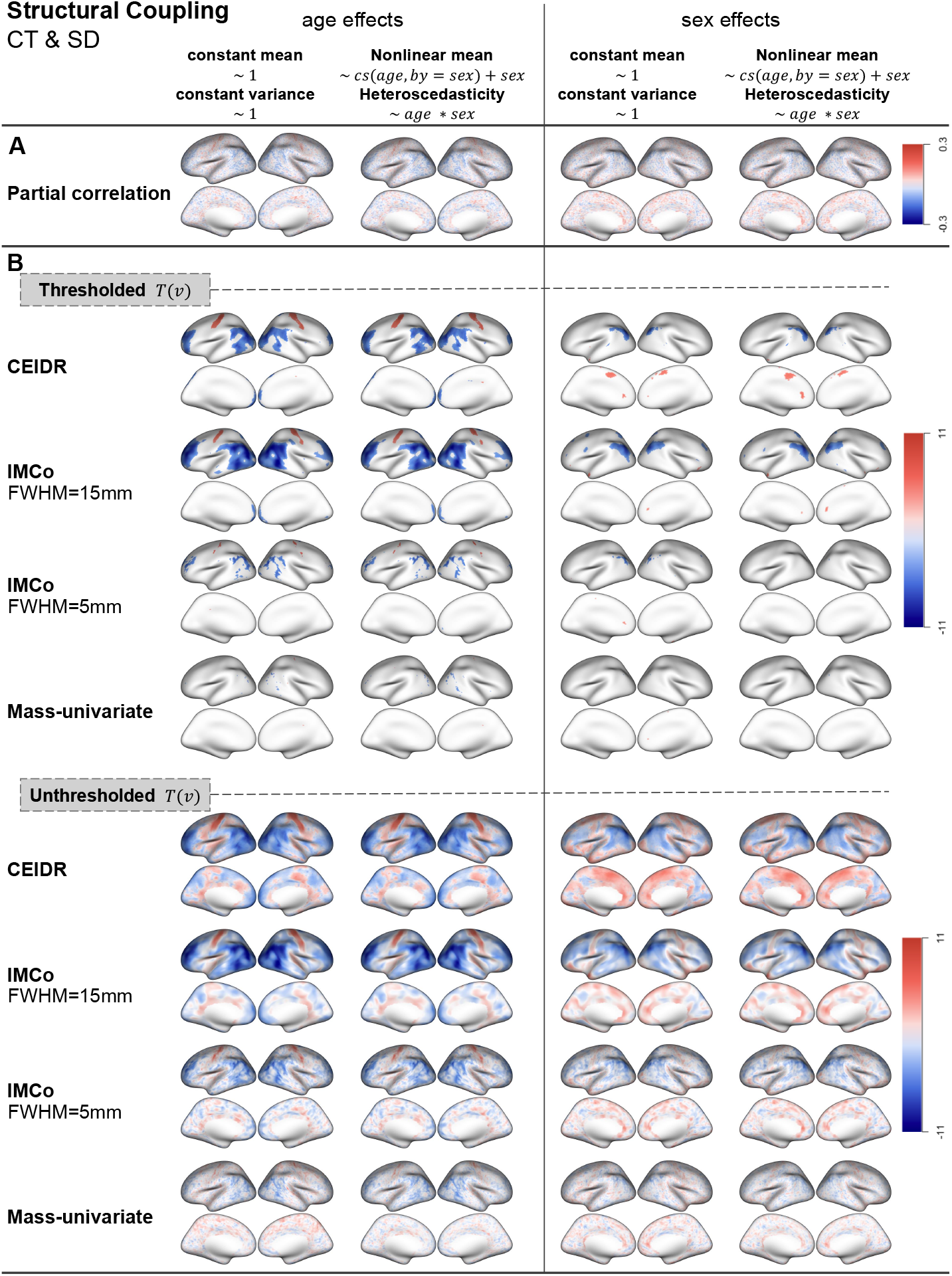
Full-sample results for CT and SD. (A) effect size maps, represented as a partial correlation of 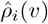 and *x*_*i*_ (e.g., age) given **z**_*i*_ (e.g., sex). (B) thresholded and unthresholded test statistic maps. Negative values of test statistics or partial correlation (blue) imply that intermodal coupling measures are found to be negatively associated with a covariate of interest, while positive values imply that intermodal coupling measures are positively associated. Note that sex is coded as a binary variable with 1 for females and 0 for males; therefore, regions where we observe positive values for sex effects would indicate where coupling was found to be higher in females than in males, while regions where we observe negative values for sex effects would indicate where it is estimated to be lower for females versus males.

**Figure 4.**
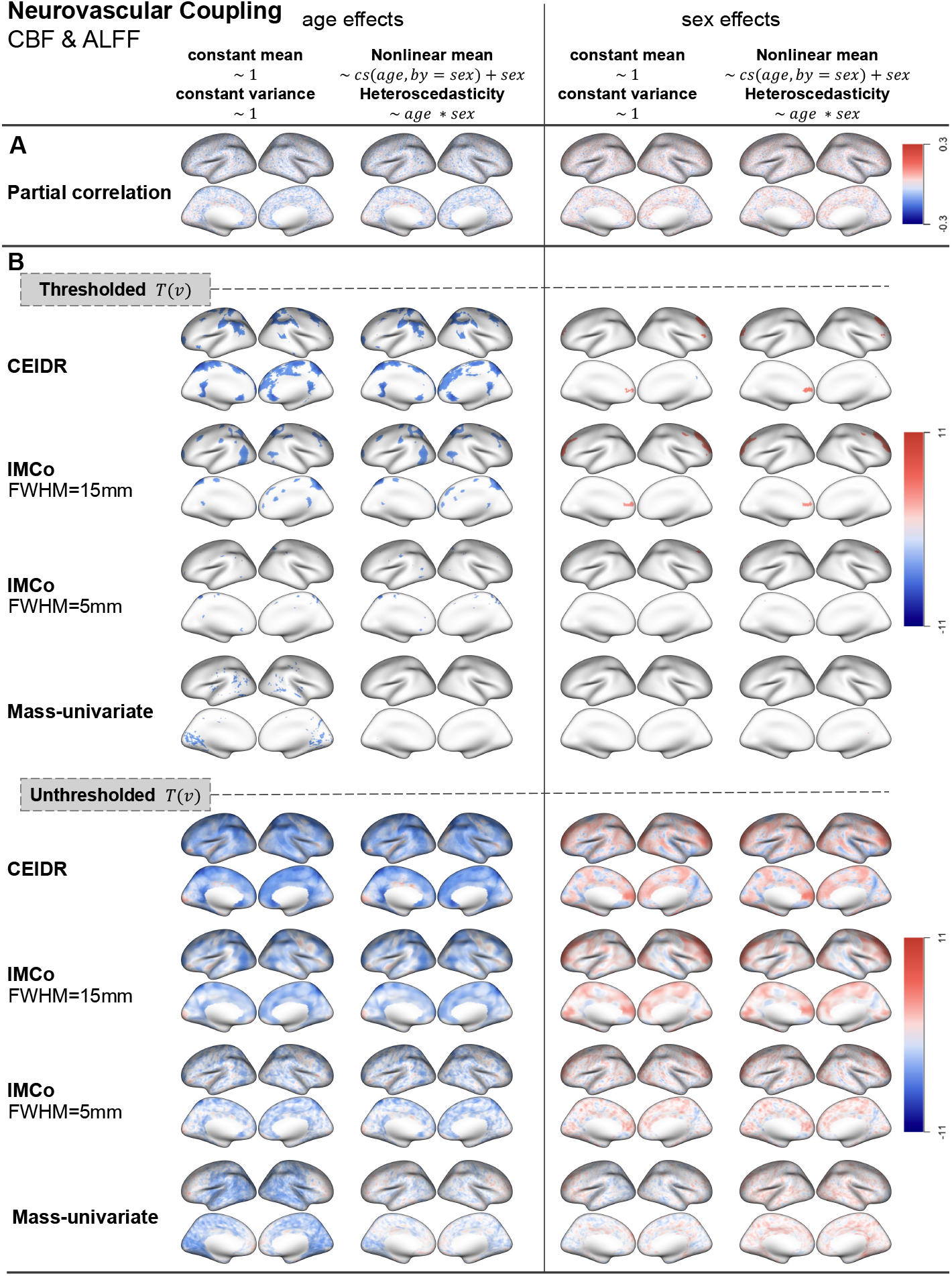
Full-sample results for ALFF and CBF, with the description analogous to Figure 3. From the formulation above, the null hypothesis *H*_0*v*_ : *κ*(*v*) = 0 can be tested by fitting the GLM and extracting the corresponding *p*-value.
IV. *Adaptive cluster enhancement*. To improve sensitivity, we perform adaptive cluster enhancement on the vertex-level score test statistics *U* (*v*) to leverage local spatial similarity of the effect sizes and improve power (Park and Fiecas 2022; Weinstein et al. 2022; Pan et al. 2024). The score test statistic at each vertex is defined as

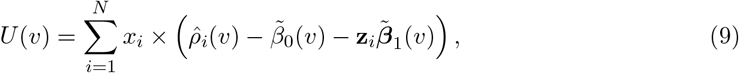

where 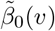,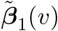 are obtained under the null model (i.e., the model without any *x*_*i*_*κ*(*v*) terms). The cluster-enhanced test statistic, denoted by *T* (*v*), is constructed by the standardized average of the vertex-level test statistics: *T* (*v*) = max*{T*_*r*_(*v*)|*r* = 0mm, 1mm,…, *r*_max_*}* where

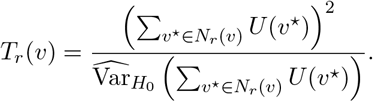
V. *Determining statistical significance*. We perform inference using a permutation-based procedure. This helps to prevent spurious associations (i.e., false positives) when working with high-dimensional neuroimaging features (Lindquist and Mejia 2015). Permutation allows us to (i) compare the observed relationship between coupling and *x*_*i*_ to observations generated from the null hypothesis, and (ii) account for multiple comparisons across all vertices by using a brain-wide threshold, *t*_*α*_, to establish statistical significance.

Because our hypothesis of interest is the covariate effect on intermodal correlations and not on means or variances in each modality, we need to permute *x*_*i*_ (or (*x*_*i*_, **z**_*i*_) when covariates are highly correlated) across subjects in stage III to generate permuted samples, while all the other steps in stages I and II are fixed. Other permutation strategies could be considered, which we refer readers to Winkler et al. (2014). A major computational advantage of this is that there is no need to redo Stage I and Stage II in each permutation, which boosts computational efficiency. Therefore, all permutation analyses can begin with the residuals 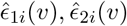. As such, this permutation scheme becomes equivalent to using the CLEAN pipeline (Park and Fiecas 2022) in stage III. For each of 1000 (or more) permutations, Stages III-IV are repeated with *x*_*i*_ or (**z**_*i*_, *x*_*i*_) permuted across subjects, which controls the family-wise error rate at a predetermined *α* (e.g., 0.05).

### 2.5 Other comparable methods

#### 2.5.1 IMCo

Vandekar et al. (2016) used locally weighted regression with pre-defined weights to estimate subject- and vertex-specific measures of intermodal coupling. That is, at each vertex, intermodal coupling was estimated by regressing one modality on the other using brain measurements. The weight is determined by (i) the distance between *v* and *v*^*⋆*^ denoted by 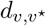 and (ii) user-specified parameter for controlling the spatial extent of neighborhood, the full width at half maximum (FWHM), with the specific form as 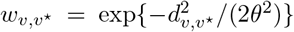 with 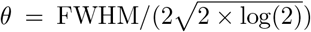. As FWHM increases, vertices that are far apart from a vertex are given more weight in fitting locally weighted regression. This procedure yields a slope at each vertex, representing local intermodal coupling for each subject. These individual-level coupling maps could then be used to test for local associations with behavioral or demographic variables.

The regression slopes derived from earlier implementations of IMCo (Vandekar et al. 2016; Baller et al. 2022) are an inherently asymmetric measure of association, since one modality must be arbitrarily chosen as the dependent variable and the other as the independent variable (Hu et al. 2022). To circumvent this discrepancy, instead of locally weighted regression slopes, we estimate locally weighted *correlations* to define subject-level intermodal coupling (Figure 2(a)).

#### 2.5.2 Mass-univariate

To emphasize the advantage of leveraging the spatial structure of the data (cluster enhancement), we also consider a “mass-univariate” approach. This involves estimating vertex-level test statistics measuring individual differences in intermodal coupling—but without leveraging neighboring information. In this article, our implementation of the mass-univariate framework is equivalent to CEIDR, when both Stage I.2 and Stage IV are excluded.

While CEIDR, IMCo, and the mass-univariate method each takes a different approach to *quantifying* subject-level intermodal coupling, we use the same permutation-based procedure for controlling FWER across the three methods. That is, given the estimate of 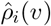 from each method for Equation (8), we randomly permute the covariate of interest, *x*_*i*_ across subjects (after adjusting for confounders **z**_*i*_), and then we repeat the steps outlined in Figure 2(b)-(d) to estimate null effects of a covariate of interest on intermodal coupling.

### 2.6 Remarks

#### 2.6.1 Comparing between-subject and within-subject adjustments

Although the between- and within-subject adjustments pursue the same goal, they rely on different assumptions and offer different advantages and limitations. The between-subject adjustment benefits from utilizing the full sample (e.g., more than 800 subjects in our real data analysis), which reduces variance of estimates. However, we note that GAM could still be misspecified and it does not address unknown covariate effects. The within-subject approach avoids reliance on covariate information, but its validity depends on the assumption that the mean and variance functions are the same within a specified neighborhood, which may be violated in practice, especially when neighbor size is large. Therefore, we recommend using a small radius to specify neighborhood (e.g., 5mm) to prevent false positives, which is used in CEIDR as a default. Although the number of vertices within a 5mm radius is small in fsaverage5 (median: 11 vertices in the pial surface), we note that adopting a higher-resolution would be helpful at the expense of increased burden in multiple comparison.

#### 2.6.2 IMCo is a special case of CEIDR

In the Supplementary Material I, we derive that IMCo can be interpreted as a special case of CEIDR under a specific setup. When equal weights are assigned to all vertices within a predefined region 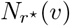 and zero weights are assigned outside this region, IMCo becomes equivalent to CEIDR with the within-subject adjustment (i.e., without between-subject adjustment) in Step I and 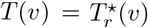 in Step IV (i.e., without *adaptive* cluster enhancement). This explains when IMCo would suffer from inflated false positives or reduced power. Specifically, when the specified full-width at half-maximum (FWHM) is large (e.g., 15mm), it would lead to biased adjustment of mean and variance (see Section 2.6.1), and power loss is expected when signal region is smaller. Similarly, when the specified FWHM is small (e.g., 5mm), false positive rate would remain appropriately controlled, but power loss is expected when signal region is larger. Therefore, the ability to separately define (i) radius used to adjust mean and variance and (ii) radii used to conduct adaptive cluster enhancement speaks to the practical utility of CEIDR.

## 3 Data analysis

### 3.1 The Philadelphia Neurodevelopmental Cohort study

The Philadelphia Neurodevelopmental Cohort is an open-source and multi-modality investigation of brain development in children aged 8 to 21 years (Satterthwaite et al. 2016). The PNC dataset includes structural neuroimaging, functional neuroimaging (resting-state and task-based fMRI), psychiatric evaluations, and a neurocognitive battery for each individual. This study is a collaboration between the Children’s Hospital of Philadelphia and the Brain Behavior Laboratory at the University of Pennsylvania, and is funded by the National Institutes of Mental Health (NIMH). Informed consent was obtained from all participating adults and guardians of underage participants, and the study protocol received approval from the Institutional Review Board at both the University of Pennsylvania and the Children’s Hospital of Philadelphia.

We apply our proposed method to multimodal neuroimaging data from the Philadelphia Neurodevelopmental Cohort (Satterthwaite et al. 2016), mapping age and sex differences in intermodal coupling between (i) cortical thickness (CT) and sulcal depth (SD) and (ii) cerebral blood flow (CBF) and amplitude of low-frequency fluctuations (ALFF). Individual differences in structural (CT-SD) and neurovascular (CBF-ALFF) coupling were also examined in previous studies involving the PNC data (Vandekar et al. 2016; Baller et al. 2022; Hu et al. 2022). Using similar exclusion criteria as in previous work, we include *N* = 911 participants in our analyses of CT-SD coupling (Weinstein et al. 2021) and *N* = 831 participants in our analyses of CBF-ALFF coupling (Baller et al. 2022; Weinstein et al. 2022).

### 3.2 Imaging parameters

Resting-state fMRI data was gathered from *N* = 1601 participants using the same scanner, a single 3T Siemens TIM Trio whole-body scanner equipped with a 32-channel head coil. Images were collected using a whole-brain, single-shot, multi-slice, gradient-echo echoplanar sequence. The sequence parameters were as follows (124 volumes): time repetition = 3000 ms, time echo = 32 ms, field of view = 192 *×* 192 mm, matrix size = 64 *×* 64, number of slices = 46, slice thickness = 3 mm, slice gap = 0 mm, flip angle = 90 degrees, and voxel resolution = 3 *×* 3 *×* 3 mm.

Brain perfusion was imaged with a 3D-encoded spin-echo pseudocontinuous arterial spin labeling sequence. The sequence parameters for perfusion were as follows (80 volumes): time repetition = 4000 ms, time echo = 15 ms, field of view = 220 *×* 220 mm, matrix size = 96 *×* 96, number of slices = 20, slice thickness = 5mm; slice gap = 1mm, and resolution 2.3 *×* 2.3 *×* 6 mm. More detailed information on these modalities and image acquisition can be found in Satterthwaite et al. 2014.

Pre-processing of BOLD signal involved field inhomogeneity correction, registration, linear and quadratic denoising, and mitigation of motion artifacts through a confound regression model, as described in Ciric et al. 2017 and Weinstein et al. 2022. The resulting BOLD signal was filtered using a first-order Butterworth filter with a pass-band of 0.01 to 0.08 Hz in order to remove trends and noise associated with physiological processes (e.g., breathing). Then, ALFF was calculated as the sum of the amplitudes of the power spectrum over all bins in the 0.01 - 0.08 Hz band. As previously described in Baller et al. 2022, CBF was calculated using parameters derived from arterial spin labelling.

The brain measurements were mapped onto the fsaverage5 pial surface, which contains 10,242 vertices in each of the left and right hemispheres. The full set of vertices is used to compute the geodesic distance matrices separately for each hemisphere. During model fitting, we exclude the medial wall vertices—888 in the left hemisphere and 881 in the right hemisphere—resulting in 9,354 vertices in the left hemisphere and 9,361 vertices in the right hemisphere for analysis.

### 3.3 Analysis results and comparisons

We compared CEIDR to IMCo (with FWHM=15mm and FWHM=5mm) and the mass-univariate approach outlined in Section 2.5. In our analysis, the permutation approach outlined in step V of CEIDR was used in all methods considered to promote fair comparisons, although previous implementations of IMCo (Vandekar et al. (2016) and Baller et al. (2022)) controlled false discovery rate (FDR). We set FWER controlled at 5%. We also made *r*_max_ = 15mm in step IV of CEIDR for a fairer comparison with IMCo with FWHM=15mm. We fitted and compared each model with two versions of Step I: (i) no adjustment made in mean and variance functions and (ii) sex-specific non-linear effects are modeled by splines and a log-linear model is applied for variance. For a single test, the total runtime for applying CEIDR on one hemisphere (e.g., left hemisphere) is around 12 minutes on MacBook Pro 2019 (2.3 GHz 8-core Intel Core i9 and 32GB RAM) without parallel computing: adjustment for mean and variance effects takes around 7 minutes, and permutation testing with 2,000 permutations takes roughly 5 minutes.

Results from our analysis of individual differences in structural (CT-SD) and neurovascular (CBF-ALFF) coupling are presented in Figures 3 and 4. Panel A of each figure shows the effect size represented by the partial correlation between 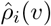 and the covariate of interest after adjusting for other covariates. In panel B, for each method, we map FWER-thresholded local test statistics *T* (*v*) quantifying individual differences in coupling. The unthresholded test statistics are also presented to aid interpretation and enhance reproducibility (Taylor et al. 2023). We will focus on interpret-ing the results for individual differences, although the estimated coupling maps are provided in Supplementary material II.

The unthresholded maps reflect similar patterns of partial correlation, larger test statistics shown in the area with higher partial correlations and smaller test statistics shown in the area with lower partial correlations. Overall, the effect size is small in both modality pairs we considered and, as a result, the mass-univariate approach yields very limited areas of statistical significance even with a relatively large sample size used in the PNC study. As expected, CEIDR and IMCo that use spatial information provide more interpretable results on individual differences in coupling, while IMCo (with FWHM=5mm) produce much less significant regions, which could be explained by the inefficiency of using such a narrow smoothing kernel in capturing broader spatial patterns in our applications. Thus, we mainly compare the results of CEIDR and IMCo with 15mm FWHM.

Maps of age effects on CT-SD coupling in Figure 3 suggest an overlap across the different implementations of CEIDR and IMCo. Both methods show patterns of negative age effects (blue) on CT-SD coupling in the visual cortex and positive age effects (red) in the motor cortex, which is consistent with the analysis conducted by Vandekar et al. (2016). Although IMCo localizes significant age effects across a larger area of the cortical surface than the other implementations, this can partially be attributed to the benefits of using a large radius (FWHM=15mm) when signal clusters are large and contiguous. In the context of CBF-ALFF coupling, however, CEIDR detects age effects across a larger portion of the cortical surface compared to IMCo. This highlights the greater effectiveness of adaptive cluster enhancement in CEIDR compared to fixed-radius enhancement in IMCo (see Section 2.6.2), particularly when signals are more scattered and form smaller clusters. Age effects appear prominent in the attention networks, which aligns with results in Baller et al. (2022).

Interestingly, qualitative differences between CEIDR and IMCo are more apparent in tests of sex effects. For CT-SD coupling, both methods detect significant negative associations between female sex and structural coupling in the parietal cortex, which is also consistent with Vandekar et al. (2016)’s results. CEIDR also reveals significant positive associations between female sex and structural coupling in the superior frontal and anterior cingulate cortices. Additionally, the two different implementations of CEIDR and IMCo consistently reveal patterns of a positive association between female sex and neurovascular coupling overlapping in the frontoparietal network, which is consistent with Baller et al. (2022).

In our analyses, no significant differences are observed between implementations using a simple between-subject adjustment model (constant mean and variance) and those employing a more complex model (nonlinear mean and heteroscedasticity). Since its effect is not clearly understood without ground truth, we next conduct a brain-level simulation study in Section 4 to evaluate false positive control and power under controlled data-generating mechanisms.

## 4 Simulation study

### 4.1 Simulation design

We conducted a simulation study to examine CEIDR’s ability to control the Type I error rate and its robustness and power. Specifically, we aim to highlight the impact of correct (vs incorrect) adjustment of marginal means and variances in mitigating false-positive rates, and to show how it relates to statistical power. As done in PNC data analysis, we considered CEIDR, IMCo (with FWHM=15mm and FWHM=5mm) and the mass-univariate approach with FWER controlled by permutation.

Data were simulated using the parametric model described in Section 2.2, with each simulated dataset consisting of *N* = 200 subjects. We first generated two covariates, 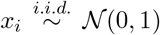 (our interest) and 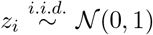 (nuisance) independently, and used them to generate brain measurements *y*_*mi*_(*v*) following Equations (1), (2), and (3). We generated various types of regions on the left hemisphere of the brain using the fsaverage5 pial surface, which is described in detail in Figure 5. These regions were designed to reflect different combinations of individual differences in marginal means, variances, and inter-modal couplings (correlations). We included two null regions, A and B, which exhibit individual differences in marginal distributions but no differences in coupling (set to *ρ*(*z*_*i*_, *x*_*i*_, *v*) = tanh(0.3 + 0.1*z*_*i*_)). Specifically, region A displayed variability across individuals in the mean and variance of modality 1, while modality 2 remained homogeneous. Region B exhibited individual differences in the means of both modalities but maintained constant variance. In addition, we defined three signal regions, C, D, and E with varying sizes, with spatial extents of 15mm, 7mm, and 5mm, respectively. In these regions, there is individual difference in intermodal correlations, which was set *ρ*(*z*_*i*_, *x*_*i*_, *v*) = tanh(0.3 + 0.1*z*_*i*_ + *κ · x*_*i*_). Region C exhibits individual differences in marginal means, variances, and coupling, whereas regions D and E exhibited individual differences in coupling only, with fixed means and variances. All remaining regions on the surface were defined as null, with constant marginal means and variances, and zero inter-modal coupling (*ρ*(*z*_*i*_, *x*_*i*_, *v*) = 0).

**Figure 5.**
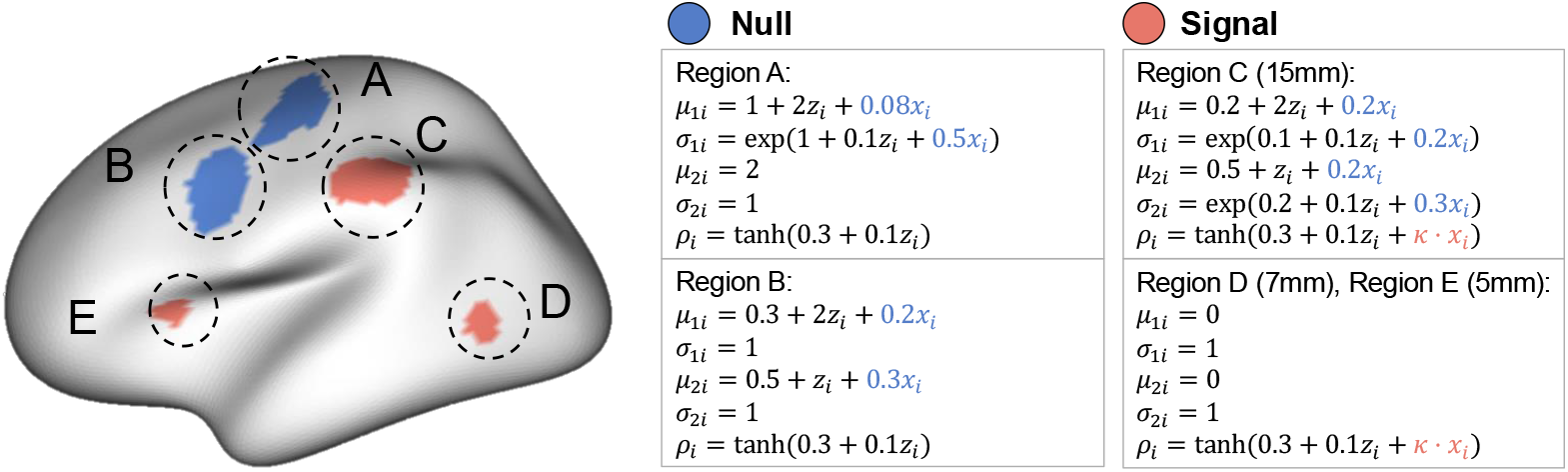
Figure for our simulation designs. Notations *µ*_*mi*_, *σ*_*mi*_, and *ρ*_*i*_ in the figure denote *µ*_*m*_(*z*_*i*_, *x*_*i*_, *v*), *σ*_*m*_(*z*_*i*_, *x*_*i*_, *v*), and *ρ*(*z*_*i*_, *x*_*i*_, *v*) respectively. Regions with blue color exhibit individual differences in means, variances, or correlations but correlations do not depend on *x*_*i*_. Regions with red color indicate the existence of the covariate effect of *x*_*i*_ in correlations when *κ* 0 regardless of the mean or variance structures. All remaining regions on the cortical surface are set as null regions with *µ*_*mi*_ = 0, *σ*_*mi*_ = 1, and *ρ*_*i*_ = 0.

In our simulations, we varied *κ* across the values 0, 0.01, …, 0.09. When *κ* = 0, regions C, D, and E reduced to null regions as well, allowing for the evaluation of family-wise error rate. When *κ≠*0, these regions exhibit nonzero covariate effects in couplings, enabling the assessment of statistical power.

### 4.2 Simulation results

Figure 6 summarizes the average performance across 1000 simulations. When *κ* = 0, the empirical FWER is defined as the proportion of simulations (out of 1000) in which at least one vertex was identified as significant. We first note that IMCo (with FWHM=15mm) had an inflated FWER. It can be explained by the presence of individual differences in means and variances in regions A, B, and C, in which IMCo’s within-subject adjustment provided biased estimates in means and variances via over-specified FWHM. In particular, outside A, B, and C exhibit different means or variances compared to the values within the regions. As a result, IMCo (with FWHM=15mm) failed to adequately adjust for these differences, particularly at the boundaries of Regions A, B, and C. In contrast, IMCo (with FWHM=5mm) still controlled the FWER, likely due to its higher weighting of smaller local neighborhoods, leading to more accurate local correlation estimates compared to FWHM=15mm. We also note that the mass univariate approach also controlled FWER accurately because it specifically modeled covariate effects using the between-subject adjustment. Altogether, these explain how CEIDR, which incorporates both between-subject and within-subject adjustments (with a small neighbor size), accurately controlled FWER.

**Figure 6.**
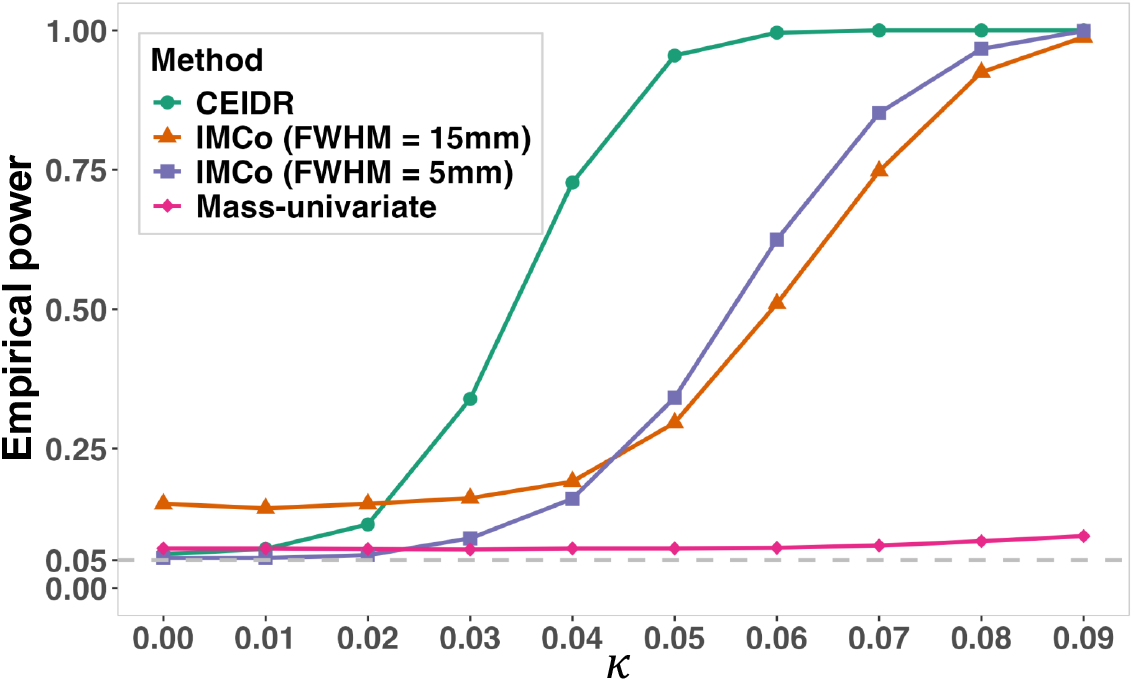
Empirical power of CEIDR, IMCo (FWHM=15mm), IMCo (FWHM=5mm), and mass-univariate from simulation study. The gray dashed line in the figure is the nominal FWER=0.05. When *κ* = 0, the corresponding value on the *y*-axis denotes empirical FWER.

As *κ* increased, the power of all three methods increased, although mass-univariate approach showed minimal increases in power as *κ* increased, which is expected. Compared to the mass-univariate approach, both CEIDR and IMCo (with FWHM=15mm and FWHM=5mm) achieved higher power through cluster enhancement. Notably, CEIDR outperformed IMCo with FWHM=5mm at all *κ* values, and even IMCo with FWHM=15mm when *κ >* 0.03. It can be explained by the varying sizes of the signal regions because the fixed clustering widths used by IMCo are either too wide (15 mm) to capture small signals effectively, or too narrow (5 mm) to capture larger signals adequately, while CEIDR adaptively captures them.

## 5 Discussion

Studies of intermodal coupling are an increasing focus of neuroscience research. Measures of intermodal coupling have previously been implicated in healthy brain development, sex differences in development, cognition, and psychopathology (Avants et al. 2010; Devonshire et al. 2012; Ouyang et al. 2015; Phillips et al. 2016; Iadecola 2017; Baum et al. 2020). However, prior methodology in this area has focused primarily on the *quantification* of subject-level intermodal coupling—and less on the power of statistical tests using these measures to establish their relevance in neurodevelopment. In this article, we sought to assess power and replicability in intermodal coupling analyses, and lay the groundwork for more rigorous and generalizable research on coupling-based biomarkers in neurodevelopmental research.

Here, we introduced a new method, called CEIDR, which may be useful in studies examining individual differences in intermodal coupling. The methodological contributions of CEIDR are twofold. First, we extend the General Linear Model (GLM) framework, commonly applied in neuroimaging data analysis, to account for conditional correlations Tu et al. (2024). In doing so, we aim to define and parametrize ‘intermodal correlation’ and its individual differences, providing a clear null hypothesis. This model-based approach enables us to mitigate potential confounding effects arising from modality-specific individual differences in means and variances that would inflate false positive findings. The family-wise error rate is rigorously controlled through permutation testing. Utilizing both structural and functional MRI data from the Philadelphia Neurodevelopmental Cohort, we identified significant age- and sex-related differences in the coupling between structural and neurovascular features in the developing brain.

In this paper, we specifically focused on analyzing neurovascular (i.e., functional) coupling between CBF and ALFF and structural coupling between CT and SD, but not structure-function coupling (e.g., CT and ALFF). While our framework is in principle applicable to structure-function analyses, our primary focus was on neurovascular coupling in the context of this neurodevelopmental cohort. We also acknowledge that several studies (e.g., Weinstein et al. (2021) and Weinstein et al. (2022)) have shown limited evidence of the ‘existence’ of coupling at the group level (e.g., *N*-back activation and CT), which might imply limited effect sizes for individual differences in their coupling. Although structure-function coupling has not been extensively explored in this dataset, there may be other contexts or modalities where such coupling is more plausible. Future applications of our framework, potentially in combination with multivariate modeling approaches (e.g., Spisak et al., 2023), may offer more insight into individual differences in structure-function coupling in modalities where such effects are expected.

In this paper, we built a methodological connection between CEIDR and IMCo from the perspective of cluster enhancement and adjustment of means and variances in each modality, and discuss some issues related to false positives. IMCo naturally depends on the user-specified FWHM and kernel function, and its spurious findings were reported in Vandekar et al. (2016) that direction (sign) of age effects on coupling change as FWHM changes. From the perspective of CEIDR, it is attributed to the risk of overspecifying FWHM in mean-variance adjustments, which could lead to false positives. On the other hand, CEIDR provides a model-based definition of coupling and their individual differences, in which evaluation of sensitivity and specificity becomes straightforward. Therefore, we believe CEIDR provides a good methodological perspective on extending intermodal coupling research at the group level.

CEIDR has several limitations, which open several opportunities for extensions of the method. In this article, we used cross-sectional data from the PNC; however, CEIDR can be naturally extended to longitudinal data, including test-retest and repeated measures analyses. Longitudinal neuroimaging studies, such as the Adolescent Brain Cognitive Development (ABCD) study, have been instrumental in advancing the understanding of adolescent neurodevelopment by allowing the modelling of individual trajectories of imaging phenotypes over time (e.g., Romer et al. (2023) and Holm et al. (2023)). Such longitudinal data would allow us to explore whether changes, and the rate of change, in intermodal coupling differ by clinical groups or sex (Park et al. 2021; Bernal-Rusiel et al. 2013a; Bernal-Rusiel et al. 2013b). Additionally, CEIDR could be extended to analyze the test-retest reliability of individual differences in intermodal coupling, an important part of validating biomarkers from brain imaging (Pan et al. 2024). Another interesting direction is to model and test non-linear age trajectories in coupling (Vandekar et al. 2016; Baller et al. 2022). Lastly, we focused on analyzing the coupling between two imaging modalities. Extending this to pairwise comparisons across multiple modalities may be critical for a more comprehensive biological understanding and could facilitate whole-brain generative modeling (Hu et al. 2022).

Another limitation of the present work is that our interpretations of localized individual differences in intermodal coupling were based on visual assessments of the brain maps presented in Figures 3 and 4. To substantiate future interpretations, we plan to integrate methods for testing spatial specificity-or “enrichment”-of brain-phenotype associations within functional networks and other regions of interest. Methods for such analyses have been recently adapted from genetics (Subramanian et al. 2005) to neuroimaging (Weinstein et al. 2024). In the context of mapping individual differences in intermodal coupling, enrichment analyses would add to the rigor and replicability of assessments of spatial specificity in multimodal neuroimaging research.

## CRediT authorship contribution statement

**RP**: Methodology, Software, Formal analysis, Validation, Visualization, Writing – original draft, Writing - review & editing. **SMW**: Conceptualization, Methodology, Software, Formal analysis, Validation, Visualization, Writing – original draft. **DT**: Conceptualization, Methodology, Validation, Visualization, Writing – original draft. **FH**: Software, Formal analysis, Writing - review & editing. **BT**: Formal analysis. **RZ**: Methodology, Writing - review & editing. **SNV**: Methodology, Validation, Writing - review & editing. **EBB**: Data curation, Software, Writing – review & editing. **RCG**: Resources, Writing – review & editing, Funding acquisition. **REG**: Resources, Writing – review & editing, Funding acquisition. **AAB**: Writing – review & editing, Funding acquisition. **TDS**: Resources, Methodology, Writing – review & editing, Funding acquisition. **JYP**: Conceptualization, Methodology, Validation, Visualization, Writing – original draft, Writing - review & editing, Funding acquisition, Supervision.

## Ethics statement

The study procedures of the Philadelphia Neurodevelopment Cohort (PNC) were approved by the Institutional Review Boards (IRBs) of the University of Pennsylvania and the Children’s Hospital of Philadelphia.

## Data and code availability

Raw neuroimaging data from the PNC study are publicly available at the dbGaP https://www.ncbi.nlm.nih.gov/projects/gap/cgi-bin/study.cgi?study_id=phs000607.v3.p2. ALFF, CT, and SD data from PNC, as well as other data types, is also publicly available for download through the Reproducible Brain Chart at https://reprobrainchart.github.io/ (Shafiei et al. 2025). All analyses were run using the R version 4.2. CEIDR is currently available for implementation at https://github.com/ruyipan/CEIDR as an R package. Runtime for applying CEIDR on one hemisphere in the PNC study (*N* = 831 subjects) is roughly 12 minutes on MacBook Pro 2019 (2.3 GHz 8-core Intel Core i9 and 32GB RAM) including permutation without parallel computing.

## Acknowledgments

We thank three reviewers for their insightful comments and suggestions, which significantly improved the quality of the manuscript. This work was supported by NIH R01MH112847 (TDS). Additional support was provided by NIH R01MH113550 (TDS), R01MH120482 (TDS), R01EB022573 (TDS), R37MH125829 (TDS), R01MH119219 (RCG, REG), K23MH133118 (EBB), R01MH123563 (SNV), R01MH132934 (AAB), R01MH133843 (AAB), and the Brain and Behavior Research Foundation NARSAD #31319 (EBB). JYP was supported by Natural Sciences and Engineering Research Council of Canada (NSERC) (RGPIN-2022-04831), the University of Toronto’s Data Science Institute (Catalyst Grant), McLaughlin Centre (Accelerator Grant), and the Connaught fund. DT contributed to this article as an employee of the University of Pennsylvania and the views expressed do not necessarily represent the views of Regeneron Pharmaceuticals Inc.

## Supplementary materials

### I. Revisiting IMCo as a special case of CEIDR

The subject-level intermodal coupling estimated by IMCo using locally weighted correlation, denoted by *ϕ*_*i*_(*v*) here, is

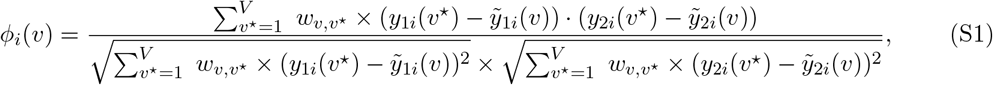

where 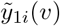 and 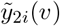 are local weighted means, computed by

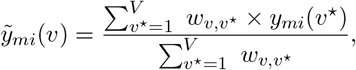

And 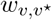 is the weight determined by the distance 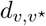 and prespecified FWHM as described in Section 2.5.1 of the paper. To build methodological connection, we consider a special case of 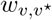 that

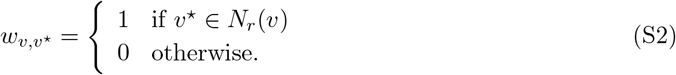

Then, 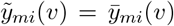 (sample mean of *y*_*mi*_(*v*) within *N*_*r*_(*v*)) defined in Section 2.4 (I.2) of the paper and Equation (S1) can be re-written as

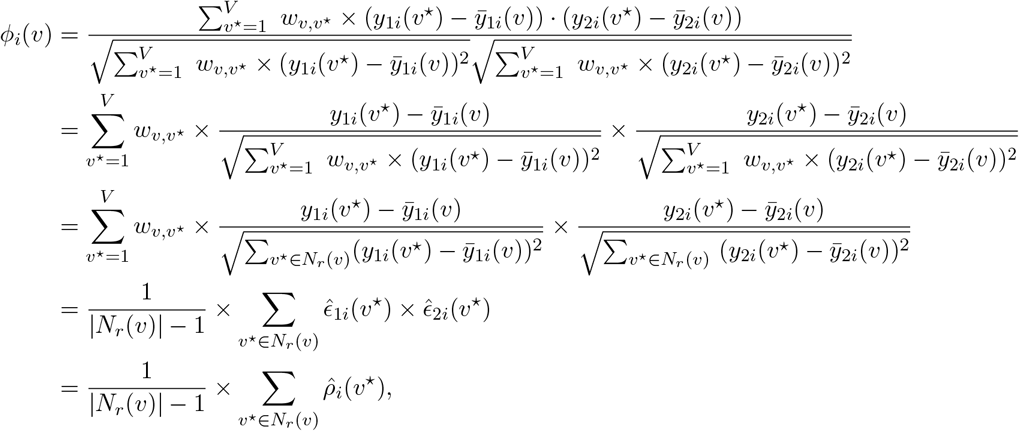

where 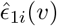 and 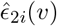 are defined in Section 2.4 (I.2) and 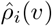 is defined in Section 2.4 (II). This result implies that IMCo’s subject-level coupling (*ϕ*_*i*_(*v*)) is proportional to the sum of CEIDR’s subject-level coupling over *N*_*r*_(*v*) after applying the within-subject adjustment using vertices of *N*_*r*_(*v*).

Once *ϕ*_*i*_(*v*) is obtained, IMCo uses GLM to compute *p* value for individual differences. Suppose that we define (i) **Z** as matrix of (1, **z**_*i*_) stacked in row across *N* subjects (ii) **x** as a column vector of *x*_*i*_, and (iii) ***ϕ***(*v*) as a column vector of *ϕ*_*i*_(*v*) collected across *N* subjects. Also, let **I**_*N*_ be a *N × N* identity matrix. Then the score test statistic for the association between *ϕ*_*i*_(*v*) and *x*_*i*_ adjusting for **z**_*i*_ is written as

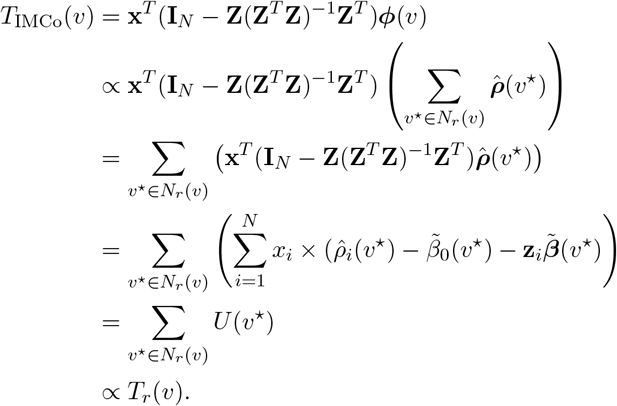

This result implies that the *p* value of IMCo at vertex *v* is equivalent to the fixed-radius cluster enhancement using score test statistics from CEIDR. When permutation to control FWER is used instead of FDR, Step V completes statistical inference done in CEIDR.

Altogether, our derivation demonstrates that, when specific weights 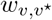 are given as Equation (S2), IMCo implemented with weighted correlation and FWER control reduces to a special case of CEIDR that (i) IMCo adjusts for means and variances in each modality by using the *within-subject adjustment* approach (Step I.2) and performs cluster enhancement with a fixed radius (Step IV).

### II. Coupling maps

In our data analysis, we computed the average intermodal coupling at each vertex by averaging 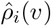 in Stage II across subjects. The resulting spatial map is presented in Figure S7.

**Figure S7.**
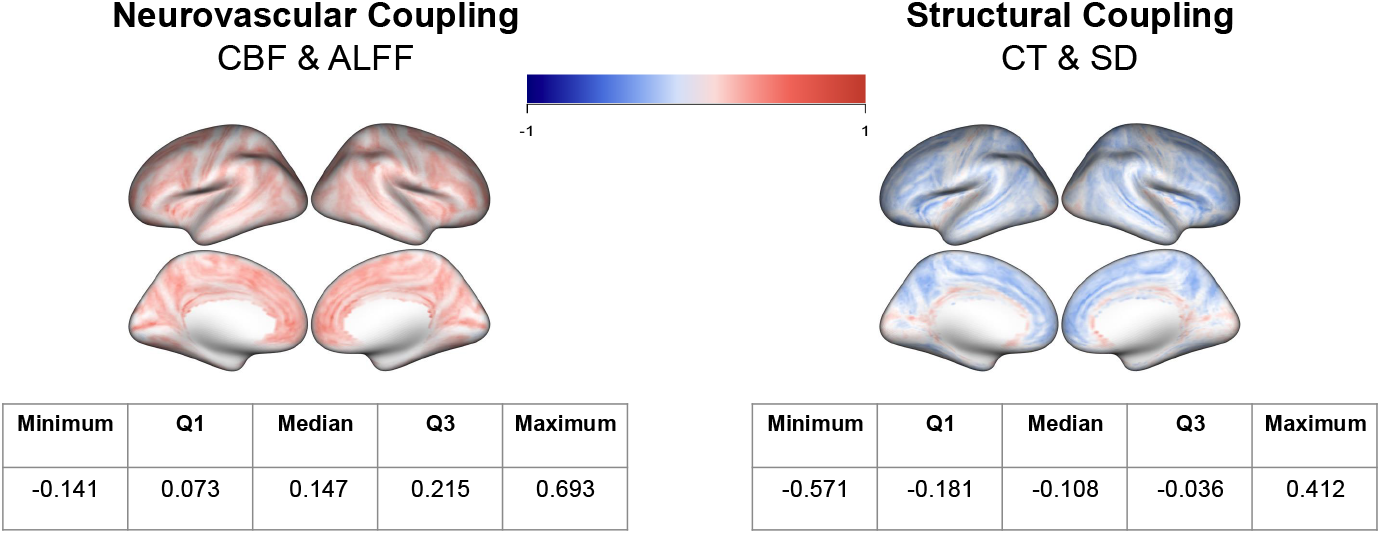
Average intermodal coupling map across subjects. For each vertex, the value represents the average of subject-specific coupling estimates computed in Stage II.

